# Exploring the beta burst dynamics of cued voluntary movements in Tourette Syndrome

**DOI:** 10.64898/2026.07.27.740916

**Authors:** Mairi S. Houlgreave, Aikaterini Gialopsou, Elena Boto, Matthew J. Brookes, Stephen R. Jackson

**Author notes:** Correspondence to: Mairi Houlgreave, School of Psychology, University of Nottingham, University Park, Nottingham, NG7 2RD, UK. E-mail address (M.S. Houlgreave).

## Abstract

Tourette Syndrome (TS) is a neurodevelopmental hyperkinetic disorder characterised by involuntary tics. These tics are thought to arise due to hyperexcitability of the motor cortex due to disinhibition within the cortico-striato-thalamo-cortical pathway. Given the involvement of motor circuits in this disorder, we explored whether there is a difference in the oscillatory dynamics of voluntary movements in people with tic disorders.

We recorded optically pumped magnetometer magnetoencephalography during cued voluntary finger abductions in individuals with tic disorders and age- and sex-matched neurotypical controls. We analysed the data both using conventional time frequency analysis and using hidden Markov modelling to explore the difference in beta burst characteristics and dynamics.

Whilst no differences were seen between groups during conventional analysis, we demonstrated an increase in beta burst duration during the post-movement beta rebound within the contralateral motor cortex in individuals with TS suggestive of increased inhibition following movement. We also show evidence of increased contralateral sensorimotor functional connectivity and reduced connectivity from and between frontal regions, as measured using coincident beta bursts. This is suggestive of disrupted functional connectivity within frontal control networks in individuals with TS.

## Introduction

Tourette Syndrome (TS) is a neurodevelopmental hyperkinetic disorder with a prevalence of ∼1% of children and adolescents worldwide (Yilmaz & Jankovic, 2025). The hallmark symptom of TS are tics; sudden, involuntary movements or sounds. The pathophysiology of TS is thought to involve striatal disinhibition within the cortico-striato-thalamo-cortical pathway giving rise to hyperexcitability of the motor cortex (Albin et al., 1989; Leckman et al., 2006, 1993).

Despite hypothesised hyperexcitability of the motor cortex, transcranial magnetic stimulation research has demonstrated a reduction in motor evoked potential (MEP) amplitude ramp up during movement preparation in TS patients, which is suggestive of cortical hypoexcitability (Draper et al., 2015; Heise et al., 2010; Jackson et al., 2013). Those TS patients who reported lower tic severity show the largest MEP amplitude increases, while those with high tic severity show a lack of MEP amplitude increases prior to a movement (Draper et al., 2015). As many patients will have a reduced severity of tic-related symptoms in adulthood, it is important to bear in mind that some of the differences seen may relate to compensation rather than pathophysiology.

Neural oscillations are rhythmic patterns of neuronal activity. When large populations of cortical neurons are synchronously active, their firing frequency becomes observable through electrophysiological techniques like EEG or MEG. During voluntary movement beta (13-30 Hz) and mu-alpha (8-12 Hz) oscillations are known to desynchronise (Pfurtscheller, 1981; Salmelin et al., 1995). Following cessation of the movement, there is an increase in beta synchronisation known as the beta rebound. A reduction in beta synchrony is thought to be an index of motor readiness (N. Jenkinson & Brown, 2011), whereas mu-alpha synchrony is associated with inhibition of brain regions not involved in the current task (Brinkman et al., 2014, 2016; Buchholz et al., 2014; Jensen & Mazaheri, 2010). MEG research by Franzkowiak and colleagues (2010), demonstrated an increase in both beta movement-related desynchronisation and rebound associated with voluntary movement in individuals with TS, but no difference in mu-alpha activity compared to controls. On the other hand, a more recent EEG study described no difference in movement related beta desynchronisation prior to voluntary movement compared with neurotypical controls, however, there was no associated mu-alpha desynchronisation (Morera Maiquez et al., 2022).

Recently, the persistent nature of neural oscillations has been questioned due to the evidence of transient bursts in activity at the single trial level in electrophysiological data (van Ede et al., 2018). This has implications on whether analysis of sustained rhythms are the best representations of the neural activity behind certain behaviours, especially as bursts can be analysed based on their duration and rate among many other measures in addition to the usual measures of power and frequency (van Ede et al., 2018). Arguably a better way to investigate changes in oscillatory activity considering these findings may be hidden Markov modelling (HMM) as classifying the timeseries data into states will provide a signal-to-noise ratio similar to that achieved by averaging over trials (van Ede et al., 2018). HMM is a statistical model which assumes the presence of hidden brain states which give rise to the observed activity within a region at each timepoint. We assume the state at each timepoint (t) only depends on the state of the timepoint before (t-1) and is therefore Markovian (Seedat et al., 2020).

Given the involvement of motor circuits in tic disorders, it is of interest whether there is a difference in the production of voluntary movements in people with tic disorders compared with neurotypical controls. If no difference is seen in the trial averaged data but differences in the burst characteristics are highlighted, this would give greater understanding of any compensatory mechanisms involved. Here, we explore the oscillatory changes within motor areas during voluntary movements in participants with tic disorders and age- and sex-matched controls using optically pumped magnetometer (OPM) MEG and HMM analysis.

## Methods

### Participants

Twenty participants with Tourette Syndrome (TS) and twenty age- and sex-matched typically developing controls were recruited. All participants were right-handed according to the Edinburgh Handedness Inventory (Oldfield, 1971). Participants gave informed consent, and the experimental paradigm received local ethics committee approval (School of Psychology, University of Nottingham). A small inconvenience allowance was provided for volunteers for their participation. Participants agreed that pre-existing structural MRI data (obtained within the Sir Peter Mansfield Imaging Centre, University of Nottingham) could be used by the researchers. Of the original 20 TS datasets collected (aged 32.7 ± 10.6 years (mean ± SD); 11 female), 2 were excluded due to trigger issues and 4 further datasets were excluded prior to analysis due to excessive noise during the MEG recording. This resulted in 14 usable TS datasets (aged 26.7 ± 3.6 years (mean ± SD); 7 female) and 20 control datasets (aged 32.2 ± 7.7 years (mean ± SD); 11 female). There was no significant difference in age or sex of participants included in the recruitment and analysis stages.

### Clinical Measures

Individuals with TS completed questionnaires, including the Yale Global Tic Severity Scale (YGTSS; (Leckman et al., 1989)), Premonitory Urge for Tics Scale – Revised (PUTS-R; (Baumung et al., 2021)) and Yale–Brown Obsessive–Compulsive Scale (Y-BOCS; (Goodman et al., 1989)), to assess their symptoms.

The YGTSS is a semi-structured interview which evaluates the number, frequency, intensity, complexity and interference of the participant’s motor and phonic tics over the previous week (Leckman et al., 1989). The PUTS-R evaluates the participants’ agreement with statements about common feelings prior to a tic using a 5-point Likert scale (0 = “Not at all”, 4 = “Very much”) (Baumung et al., 2021). PUTS severity scores were calculated using items 16-24 with item 23 excluded due to its bidirectionality. Items 17 and 21 were reverse-coded. The Y-BOCS assesses the time occupied, interference, distress, ability to resist and degree of control for obsessions and compulsions separately using a 5-point Likert scale (0 = “No symptoms”, 4 = “Extreme symptoms”) (Goodman et al., 1989). Participant diagnoses, medication status and symptom severities can be seen in Table 1.

**Table 1.**
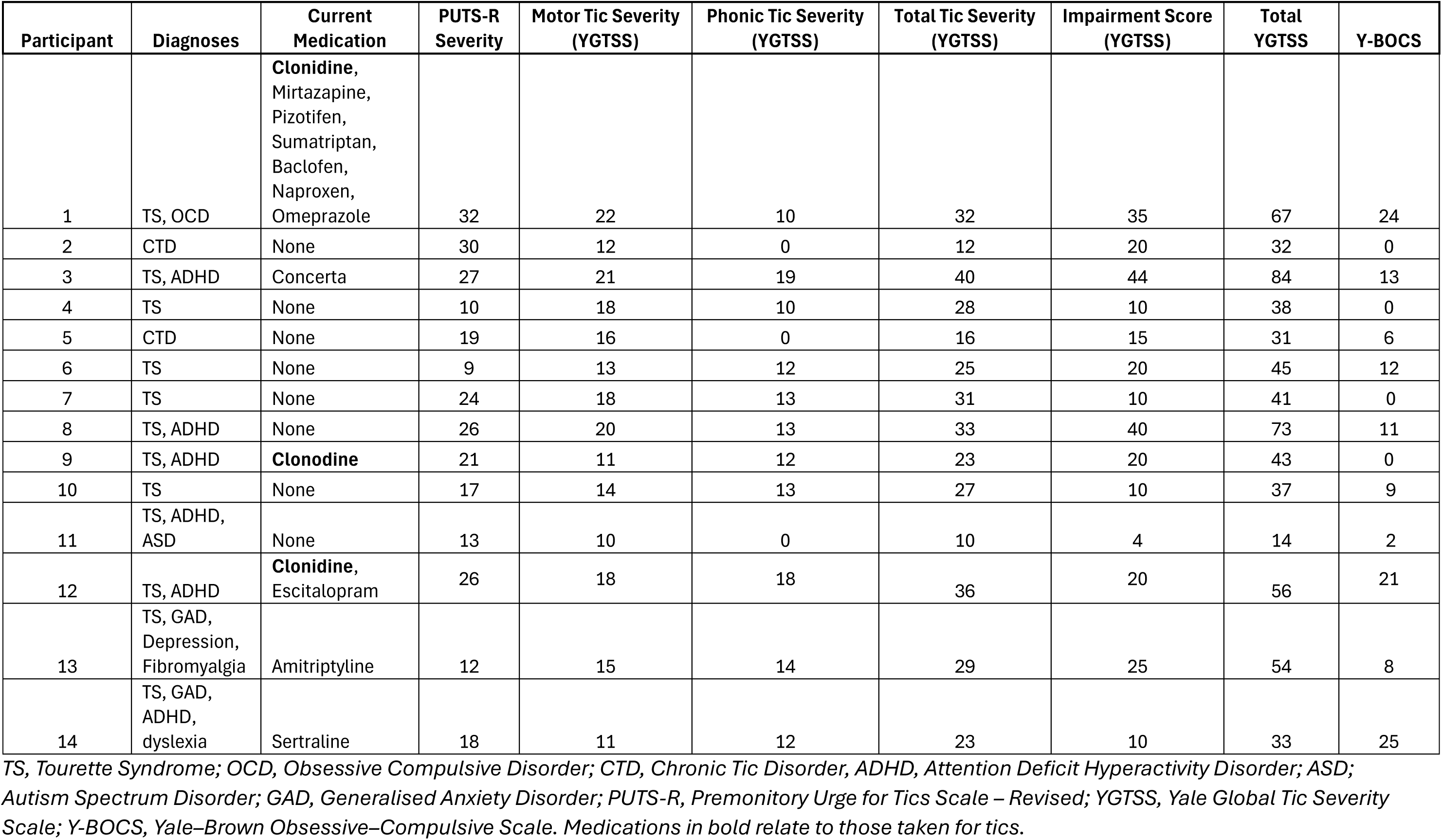
Tic Disorder participant characteristics.

### Task

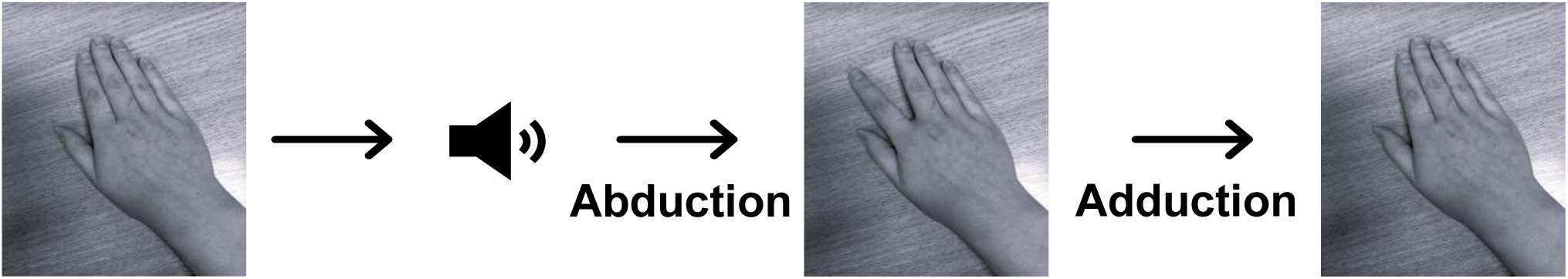

All participants completed 60 trials of a voluntary movement task where they were instructed to complete a single right index finger abduction followed by an adduction in response to an auditory cue (every 10 seconds). For participants with TS, video recordings were collected to determine whether tics interrupted these movements. If tics had interrupted the movement in a trial, then this trial would be removed from the analysis.

### Data collection

OPM-MEG data were collected at University of Nottingham using 64 triaxial OPMs (192 channels) (QuSpin Inc., Boulder, CO, USA) in a rigid adult helmet (Cerca Magnetics Ltd., Nottingham, UK) providing whole-head coverage. Participants were in a seated position. Prior to data collection the magnetically shielded room was degaussed and the remnant field was nulled. MEG data were sampled at 1200 Hz.

Following the scan, 3D head digitisations were collected both with and without the OPM helmet (Einscan H, SHINING 3D, Hangzhou, China). The head digitisation without the helmet was used for co-registration to the participants T1-weighted MRI using Meshlab (2022.02 (Cignoni et al., 2008)). Sensor location was determined through co-registration of the digitisation including the helmet.

### Pre-Processing

The data were first filtered between 1 and 48 Hz. Following this, the data were segmented such that a trial started 2s before the auditory cue for movement and ended 8s after the cue. Trials were 10s long in total. Data were inspected visually in MATLAB and any noisy/bad channels or trials were removed. Significantly more trials were included in analysis for the HC group compared to the TS group total (number of trials = 55 ± 9 (HC), 36 ± 9 (TS) (mean ± SD)) (two sample T-Test, p < 0.001). Of the 192 channels, on average 37 ± 26 were ‘bad’ for the TS group and 40 ± 10 were ‘bad’ for the HC group. There was no difference between the groups.

### Source Localisation

A linear-constrained minimum-variance (LCMV) beamformer was applied to the pre-processed data (Van Veen et al., 1997). A 4mm MNI (Montreal Neurological Institute) template brain was warped with respect to the subject’s downsampled (4 mm) anatomical scan, using FLIRT (FMRIB’s Linear Image Registration Tool) (M. Jenkinson et al., 2012). The same transformation was applied to the AAL (Automated Anatomical Labelling) atlas (Tzourio-Mazoyer et al., 2002). This resulted in an individualised parcellated cortex (78 regions) (Gong et al., 2009). Covariance was calculated for the entire experimental time window within a 1–48 Hz frequency window. The covariance matrix of the filtered data was regularised using the Tikhonov method, with the regularisation parameter set at 5% of the maximum singular value. The forward model was computed using dipole approximation and the forward model was computed using a single shell (Nolte, 2003), created using FieldTrip (Oostenveld et al., 2011). Beamformer weights were calculated for all voxels. Beamformer weights were applied to both the filtered (1-48 Hz) and unfiltered OPM data and these timecourses were then weighted towards the centroid of each brain region resulting in 78 timecourses (Brookes et al., 2016). A symmetric orthogonalisation method was used for spatial leakage correction (Colclough et al., 2015).

### Time Frequency Analysis

To statistically analyse the difference in the mu-alpha and beta timecourses between groups, the unfiltered regional timecourse was filtered into the frequency band of interest (8-12 Hz or 13–30 Hz). The absolute value of the Hilbert transform (HT) gave the magnitude of the signal at each point in time for the frequency band of interest. Each frequency timecourse was standardised (Z-scored) before being averaged across trials. The resulting timecourses were then averaged across subjects.

### Hidden Markov Modelling

To investigate differences in beta bursts which may have been averaged out when exploring trial effects, we used a 3-state Time-Delay Embedded HMM (Seedat et al., 2020; Vidaurre et al., 2018) where each state was associated with a different autocovariance pattern (time window: 230ms). The method has previously been described in Seedat et al., (2020). Briefly, the filtered (1-48 Hz) timecourse (downsampled to 100 Hz) from each AAL region was modelled independently. For each region, the state which showed maximal correlation between its probability timecourse and the amplitude envelope of the beta oscillations was taken as the ‘beta burst state’ (Seedat et al., 2020). A burst was assumed to have begun at the timepoint when the state probability surpassed a threshold of 2/3.

Several characteristics can be taken from these burst states including burst amplitude, the maximum value of the beta envelope within a burst; burst count, number of visits to the burst state (normalised by time); burst duration, time spent in the burst state.

### Functional Connectivity

Coincident bursts were used to explore functional connectivity. For all combination pairs of AAL regions, the similarity between the binary burst timecourses was calculated using the Jaccard index (Seedat et al., 2020). The larger the Jaccard index for a pair of regions, the more coincident the bursts and the higher the functional connectivity. The resulting matrices were averaged across participants and converted into pseudo-z-statistics. Global connectivity was calculated using the mean Jaccard index for the upper triangle of the connectivity matrix excluding the diagonal to ensure each pair of regions was only included once.

### Statistical Analysis

When comparing timecourses from the conventional analysis approach and for burst characteristics between groups a cluster-based permutation test (1000 permutations) was performed to control for multiple comparisons when using Wilcoxon rank sum tests.

For exploring the relationship between burst characteristics/ power spectral density (PSD) and clinical variables Spearman’s rank correlations were used. When correlating burst characteristics (probability, amplitude, duration) during desync (0-1s relative to auditory cue) and rebound (1-4s relative to auditory cue) periods with YGTSS total tic severity, YGTSS motor tic severity and PUTS severity scores, the threshold was set at p = 0.0028 to account for the 18 tests used (3 characteristics x 2 time windows x 3 clinical scores) (Bonferroni correction for multiple comparisons). This was done for the burst characteristics for the contralateral and ipsilateral motor cortices separately. Had any correlations survived Bonferonni correction a partial correlation analysis would have been performed to explore the influence of Y-BOCS scores, ADHD diagnosis and medication.

## Results

### Comparison of Trial Averaged Timecourses

There were no significant differences in the mu-alpha timecourse for participants with TS and matched neurotypical controls when analysed using a conventional time frequency analysis approach (p > 0.05, corrected) (Figure 2A). One second after the cue to move there was a brief period where the beta timecourses were significantly higher in participants with TS (p ≤ 0.05, corrected) (Figure 2B). No significant differences were seen in the ipsilateral cortex for mu-alpha and beta (Figure 3).

**Figure 2.**
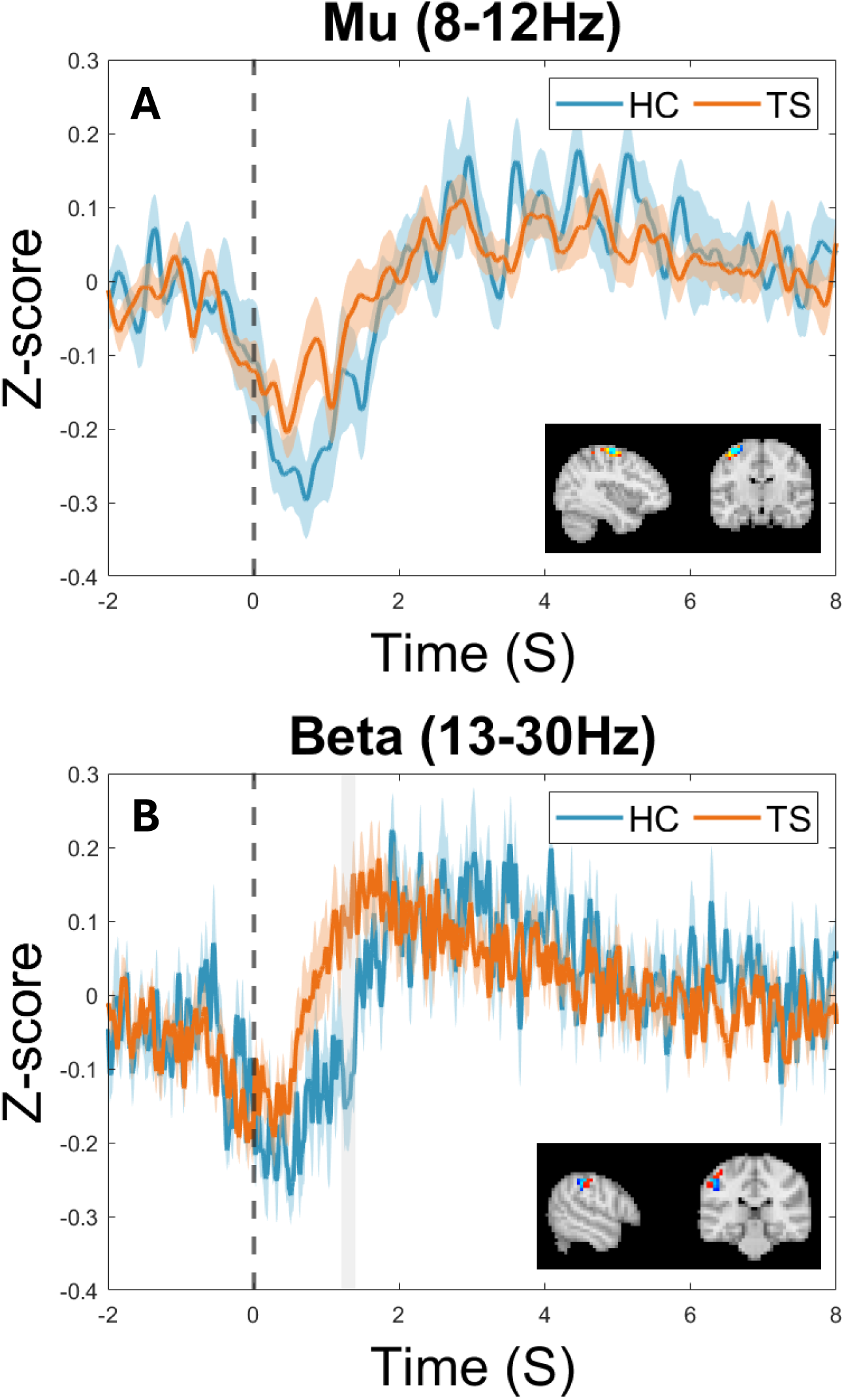
Trial averaged timecourses for (A) mu-alpha (8-12 Hz) and (B) beta (13-30 Hz) band oscillations within the contralateral primary motor cortex during and following a single index finger abduction and adduction at time 0s. Areas shaded in grey indicate a significant difference between the groups (p ≤ 0.05, corrected for multiple comparisons). In the bottom right corner of each graph there is an image of the location of the peak desynchronisation for each group TS (orange), HC (blue).

**Figure 3.**
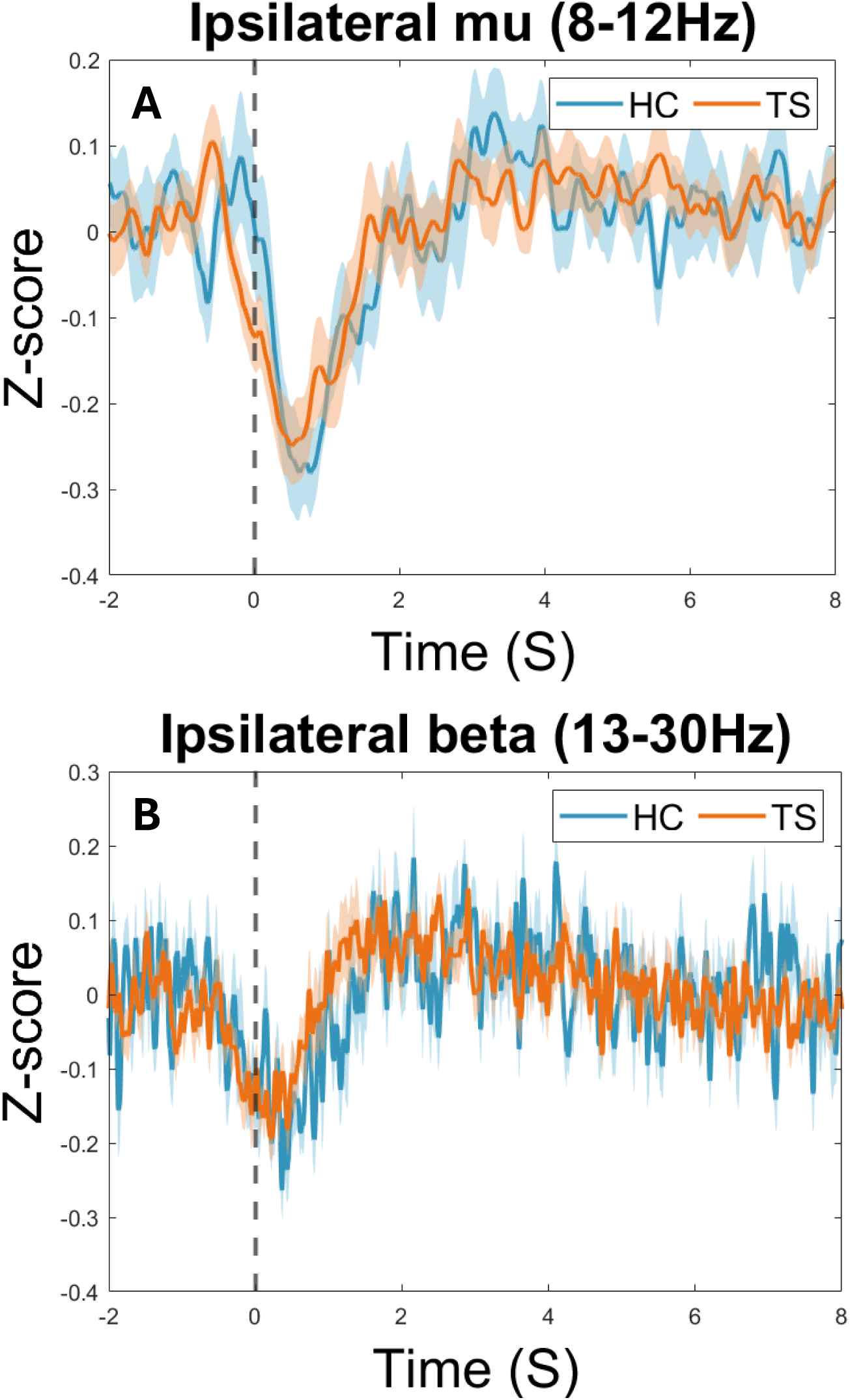
Trial averaged timecourses for (A) mu-alpha (8-12 Hz) and (B) beta (13-30 Hz) band oscillations within the ipsilateral primary motor cortex during and following a single index finger abduction and adduction at time 0s (no significant differences, p > 0.05, corrected).

### Comparison of Contralateral Burst Dynamics

Comparison of the PSD of the beta burst state within the contralateral primary motor cortex in the TS group compared to the typically developing controls showed no significant differences in the delta (p= 0.112), theta (p = 0.148), mu-alpha (p = 0.341) or beta (p = 0.2174) bands (Figure 4A).

**Figure 4.**
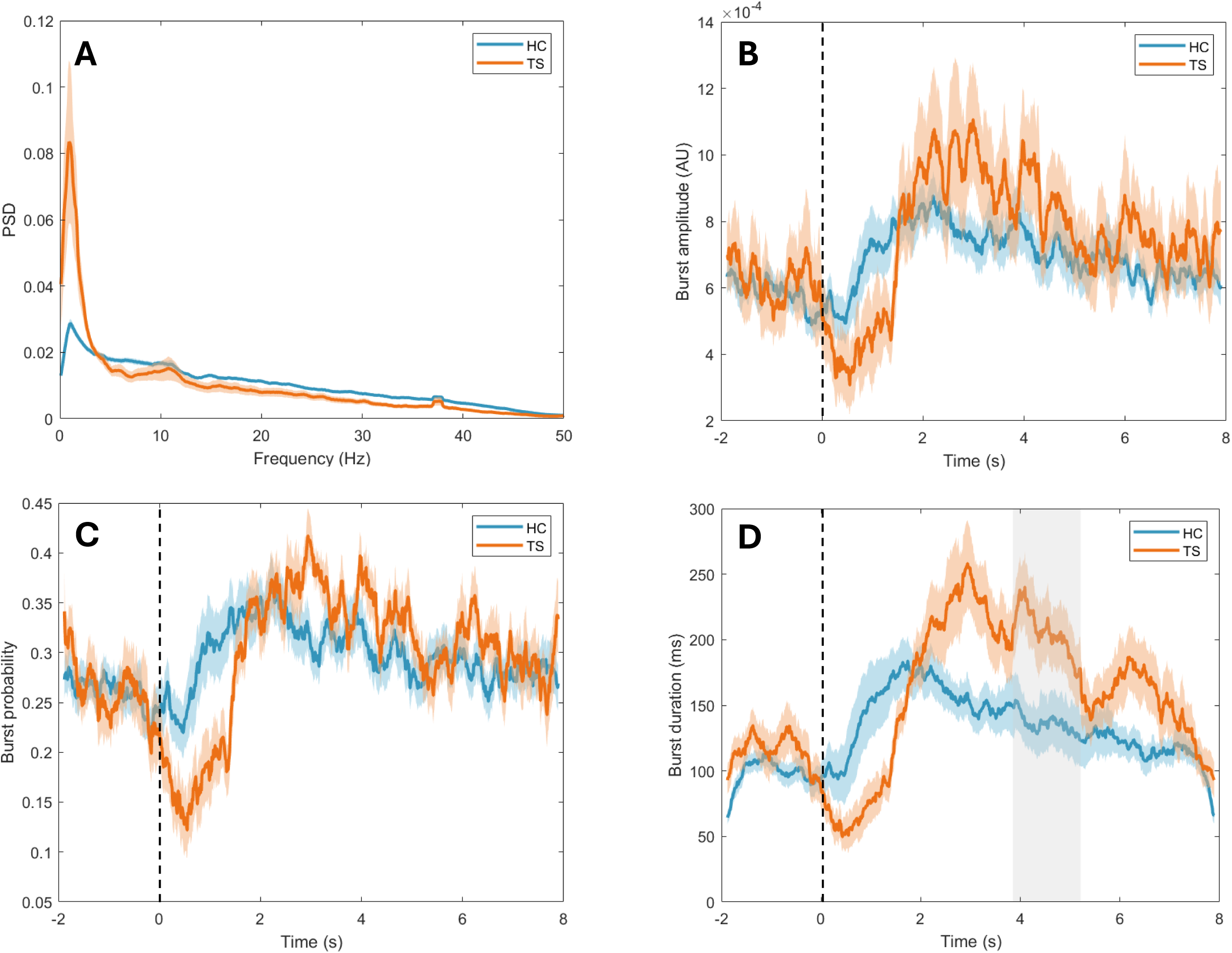
Trial averaged timecourses for characteristics of beta bursts within the contralateral primary motor cortex in individuals with Tourette Syndrome (orange) and neurotypical age- and sex-matched controls (blue). (A) power spectral density (PSD), (B) burst amplitude, (C) burst probability and (D) burst duration. A single index finger abduction and adduction occurred at time 0s. Areas shaded in grey indicate a significant difference between the groups (p ≤ 0.05, corrected).

There was evidence of a reduction in burst probability during movement and an increase in burst probability and amplitude following movement in the TS group compared to the neurotypical controls, although this did not survive multiple comparison correction (p > 0.05, corrected) (Figure 4B, 4C). There was a significant increase in burst duration during the post-movement beta rebound in the TS group compared to the neurotypical controls (p ≤ 0.05, corrected) (Figure 4D).

### Comparison of Ipsilateral Burst Dynamics

Comparison of the PSD of the beta burst state within the ipsilateral primary motor cortex in the TS group compared to the typically developing controls showed a trend for a difference in theta (p = 0.0512), mu-alpha (p = 0.077) and beta (p = 0.077) bands and no significant differences in delta (p= 0.436), (Figure 5A).

**Figure 5.**
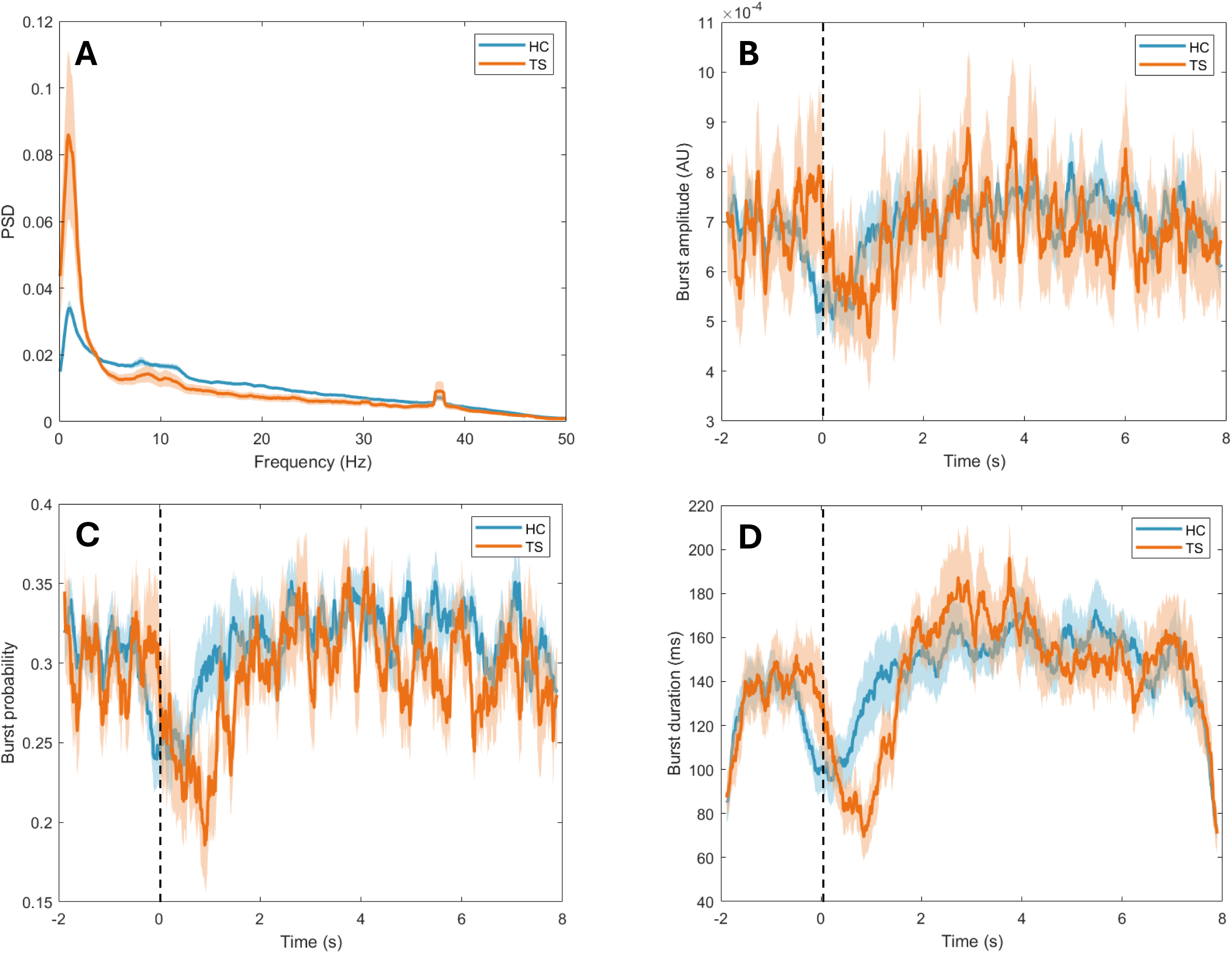
Trial averaged timecourses for characteristics of beta bursts within the ipsilateral motor cortex in individuals with Tourette Syndrome (orange) and neurotypical age- and sex-matched controls (blue). (A) power spectral density (PSD), (B) burst amplitude, (C) burst probability and (D) burst duration. A single index finger abduction and adduction occurred at time 0s (no significant differences, p > 0.05, corrected).

There was no significant difference in the timecourse of beta burst amplitude, probability or duration changes within the ipsilateral primary motor cortex during movement between the groups (p > 0.05, corrected) (Figure 5). Although there was a trend for reduced burst probability and duration during movement which did not survive multiple comparison correction.

### Comparison of Functional Connectivity

Participants with TS showed significant connectivity between left frontal regions and between occipital regions (Figure 6). There was also significant connectivity within a sensorimotor network comprising the bilateral primary sensory cortices, primary motor cortices, supplementary motor cortex (SMA), cingulum and the left insula. Similarly, matched neurotypical controls showed significant connectivity between bilateral frontal areas with a left dominance and within right sensorimotor areas (Figure 6). There was increased occipital and left sensorimotor network connectivity in people with TS compared to neurotypical controls, whereas frontal connections were weaker (Figure 7). There was however no significant difference in global connectivity between the two groups.

**Figure 6.**
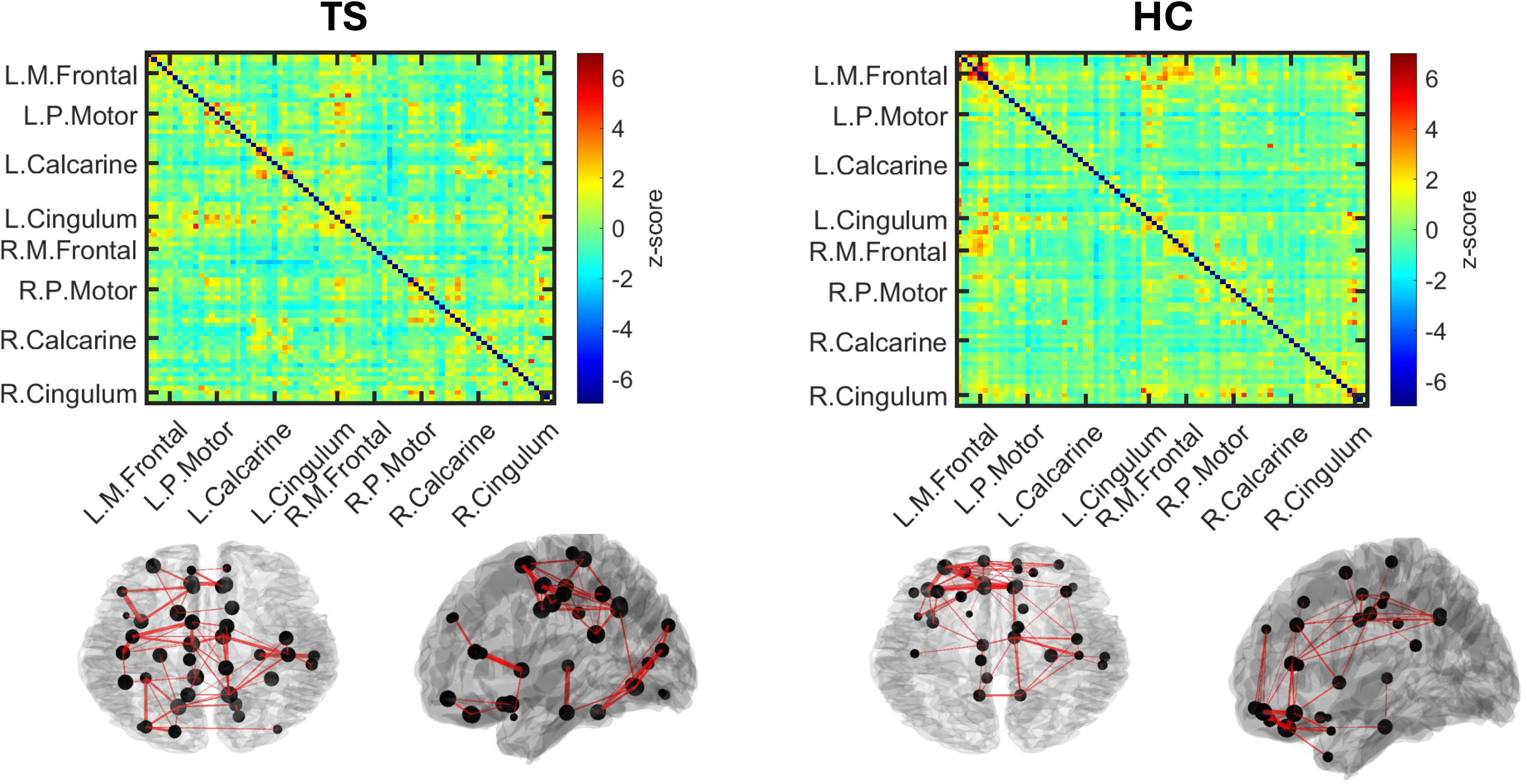
Coincident bursts in individuals with Tourette Syndrome and neurotypical age- and sex-matched controls. Jaccard matrices show all-to-all connections while the associated glass brain plots show connections where z > 2.3.

**Figure 7.**
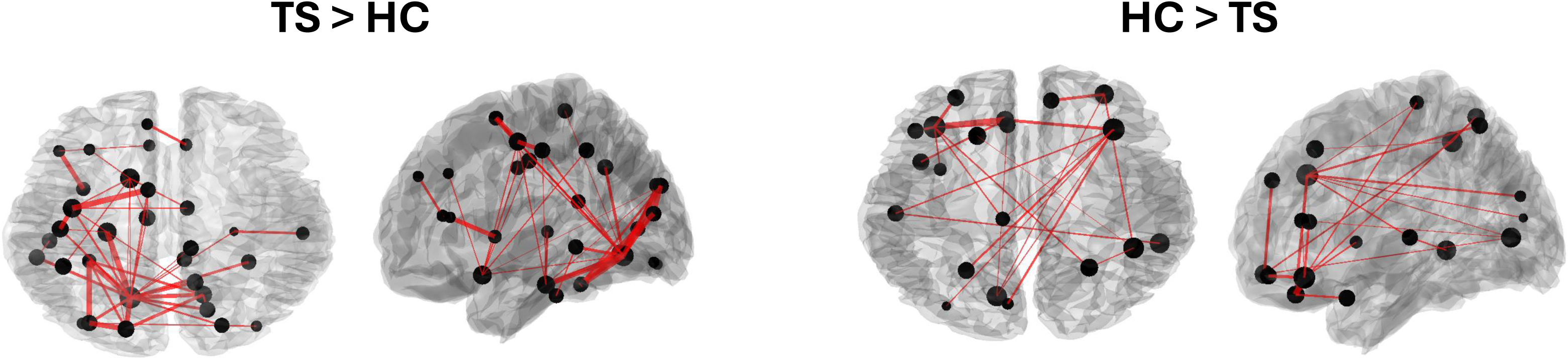
Contrasting coincident bursts in individuals with Tourette Syndrome (TS) and neurotypical age- and sex-matched controls (HC). Glass brain plots show connections where z > 2.3 for the difference between the TS and HC groups.

### Correlation with clinical measures

There was no significant relationship between total tic severity (YGTSS), motor tic severity (YGTSS) or PUTS severity scores and global connectivity (p > 0.05). Furthermore, there were no significant associations between total tic severity (YGTSS), motor tic severity (YGTSS) or PUTS severity scores and the peak in desynchronisation and rebound for burst characteristics (probability, amplitude, duration) within the contralateral motor cortex or ipsilateral motor cortex (p > 0.05). Although, there was a trend for an inverse relationship between the peak maximum burst amplitude in the contralateral (p = 0.100, R = -0.4575 (uncorrected)) and ipsilateral motor cortex (p= 0.0648, R = -0.5061 (uncorrected)) during the post-movement beta rebound and motor tic severity.

While there were no significant associations between total tic severity (YGTSS), motor tic severity (YGTSS) or PUTS severity scores and area under the curve for any frequencies within the PSD plots for contralateral or ipsilateral beta bursts were significant. There was a trend for a positive relationship between delta and motor tic severity (contralateral: p= 0.0883, R= 0.4720; ipsilateral: p= 0.0679, R= 0.5012 (uncorrected)). There was also a trend for an inverse relationship between beta and motor tic severity (contralateral: p= 0.0751, R= -0.4903; ipsilateral: p=0.0442, R= - 0.5442 (uncorrected)).

## Discussion

We investigated the difference in oscillatory dynamics during cued voluntary movements in participants with TS compared to age- and sex-matched neurotypical controls. Our results show that differences between the groups cannot be seen when analysing mu-alpha frequency timecourses using a conventional averaging approach. A brief period of difference was seen during the beta frequency timecourse, suggesting a faster recovery from movement-related desynchronisation (MRD) in the TS group. Using HMM to look at the characteristics of beta bursts over the timecourse of movement demonstrates a significantly longer burst duration during the post-movement beta rebound (PMBR) within the contralateral cortex in the TS group. This is suggestive of increased inhibition following voluntary movements. Furthermore, we see differences in functional connectivity where individuals with TS have weaker connectivity with frontal regions but increased connectivity with contralateral sensorimotor and occipital regions.

### Increased inhibition following movement

Previous research exploring differences in oscillatory activity over the contralateral motor areas prior to self-paced voluntary movements describes a significantly larger beta MRD and a trend for increased PMBR in individuals with TS compared with age- and gender-matched controls (Franzkowiak et al., 2010). A significantly higher beta amplitude was also seen during the PMBR period within the ipsilateral cortex. Our inability to replicate the increased MRD and PMBR within our conventional analysis may relate to our broadband approach. Our timecourses focus on the 13-30 Hz range, whilst the previous study by Franzkowiak and colleagues looked at the 2 Hz either side of each individual’s peak beta frequency. Another research study analysed using a similar broadband approach also found no difference in beta power between groups during cued responses (Morera Maiquez et al., 2022). However, we do report a significant increase in burst duration during the PMBR period. These results suggest that the trend for increased inhibition following voluntary movements theorised by Franzkowiak and colleagues is due to this prolonging of beta bursts, as well as potentially increased probability and amplitude as indicated by the trends in our data. We do also see increased MRD as evidenced by a lower burst probability, duration and amplitude during movement, however these differences did not survive multiple comparisons correction.

The current finding of no significant difference in mu-alpha timecourses between the groups replicates previous work looking at self-paced volitional movements but this contrasts with Morera and colleagues (2022) who found a lack of mu-alpha desynchronisation during cued ‘Go’ responses in individuals with TS (Franzkowiak et al., 2010; Morera Maiquez et al., 2022). This difference may relate to the task involved. Participants with TS have been shown to have decreased fMRI BOLD activation within the contralateral sensorimotor cortex during ‘Go’ trials compared to neurotypical controls during a Go/No-Go task (Thomalla et al., 2014). This is theorised to be related to compensatory control mechanisms in the TS group. This type of control was not required for the cued movement task described here.

Given that there is no difference in the trial averaged beta and mu-alpha timecourses, but there is a trend for increased MRD and an increase in PMBR in our beta burst data we hypothesise that this may represent a compensatory mechanism to maintain performance in the motor task and suppress tics. When at rest individuals with tic disorders are thought to display an excitation: inhibition imbalance whereby inhibition is reduced (Heise et al., 2010). However, when engaged in a goal-driven activity tics are known to reduce (Misirlisoy et al., 2014). Therefore, it would be of interest to explore beta characteristics while individuals are free to tic and whilst they are suppressing.

### Relationship with clinical measures

While none of the correlations between clinical measures survived multiple comparisons correction there was trend suggesting there may be an inverse relationship between burst amplitude during the PMBR period within the contralateral and ipsilateral cortices and motor tic severity. A similar anticorrelation between the amplitude of ipsilateral event-related synchronisation and tic severity was seen in Franzkowiak et al., 2010. The authors suggested that the ipsilateral cortex may contribute to tic suppression during volitional movement (Franzkowiak et al., 2010).

### Differences in functional connectivity

Individuals with TS show immature connectivity within the fronto-parietal and cingulo-opercular control networks when compared with age-matched controls (Church et al., 2009). This is suggestive of a delay in the development of functional connectivity in individuals with TS leading to less efficient communication within control networks. This may be reflected in our findings of reduced functional connectivity from and between frontal regions.

Our finding of increased contralateral sensorimotor network connectivity replicates previous findings of increased contralateral primary motor cortex and SMA connectivity during a self-paced movement task (Franzkowiak et al., 2012). This was hypothesised to be a marker of pathophysiology due to there being no relationship with measures of tic severity (Franzkowiak et al., 2012). Here we looked at whole brain functional connectivity, and as this increase is localised only to the contralateral side, we hypothesise that this increase in connectivity may either be a compensatory mechanism or a state-dependent marker of pathophysiology.

A previous resting-state fMRI study described an increase in local connections within the occipital lobe, similar to what is described in this study (Shprecher et al., 2014). We hypothesise that this increase in connectivity relates to the high prevalence of ocular tics in this population. In a cued blink task, children with chronic tic disorder showed highest dipole density in posterior regions whereas neurotypical controls showed highest dipole density in frontal regions (Loo et al., 2019). After the cue to blink an increase in information flow was found from occipital to frontal regions in the TS group.

### Limitations

Increased power within delta frequency band as shown in the beta burst PSD could be due to increased movement in the TS group. To ensure the differences we see in burst characteristics were not due to this we filtered the data between 5 and 48Hz and repeated the HMM analysis. Supplementary figure 1 demonstrates we replicated our findings suggesting that the differences in beta burst characteristics were not driven by high low frequency power in the TS group.

Participants with comorbidities and those taking medications for their tics were included in the TS group. As 90% of individuals with TS have a comorbid diagnosis, inclusion of those with comorbidities is more representative of the TS population. Had any significant relationships between clinical scores and burst characteristics been identified, comorbid diagnoses, Y-BOCS scores and medication status would have been included as factors to determine whether the differences survived. TS is more commonly diagnosed in males compared to females, however here we have an even split of sexes in our sample. As females with TS have a higher likelihood of severe tics enduring into adulthood and report a larger impact on quality of life (Garris & Quigg, 2021; Lewin et al., 2012; Lichter & Finnegan, 2015), a more equal split in sexes is expected in a research sample of adults.

Fewer participants were included in the TS group and of those participants there were significantly fewer trials included in the analysis. This is to be expected due to the presence of tics in this group. This will have reduced statistical power and reduced the likelihood of identifying differences between the groups. This has been mitigated through the use of non-parametric statistical methods.

## Conclusion

To conclude, our study has demonstrated an increase in beta burst duration during PMBR within the contralateral motor cortex in individuals with TS suggestive of increased inhibition following movement. We hypothesise these changes in beta burst characteristics to relate to compensatory changes to reduce tics during the task. Furthermore, our results indicate reduced functional connectivity from and between frontal regions. This provides further evidence of disrupted connectivity within control networks in individuals with TS. As these differences in the beta timecourses were only seen using beta burst analysis, based on non-trial averaged data, and little difference is seen when analysed in a conventional trial averaged analysis approach it warrants the use of beta burst analysis in other movement disorders to identify disrupted oscillatory activity.

## Data and Code Availability Statement

The MATLAB code for the voluntary movement task and analysis is available on OSF with an associated DOI: https://doi.org/10.17605/OSF.IO/9P6GD. As the MRI data used during analysis were not collected by the researchers involved, we do not have ethical approval to share these. Anonymised beamformed data can be made available on request if a formal data sharing agreement is in place.

## CRediT authorship contribution statement

M.S.H.: Conceptualisation, Data curation, Formal analysis, Investigation, Methodology, Software, Visualisation, Writing—Original draft.

A.G.: Investigation.

E.B.: Conceptualisation, Methodology.

M.J.B.: Conceptualisation, Methodology, Resources, Writing - Review & Editing, Supervision.

S.R.J: Conceptualisation, Methodology, Writing - Review & Editing, Supervision, Funding acquisition.

## Acknowledgements

Data collection for this paper was supported by research grants from NIHR Nottingham Biomedical Research Centre and Medical Research Council (MRC) (T032588). Stephen Jackson and Mairi Houlgreave were funded by NIHR-funded Nottingham Biomedical Research Centre. Stephen Jackson and Aikaterini Gialopsou were funded by MRC Grant (T032588). The views expressed are those of the authors and not necessarily those of the NHS, the NIHR, MRC, or the Department of Health. We would like to thank Caitlin Mairi Smith for sharing of participant anatomical scans for individuals with tics.

