## Supplementary Material for "Exploring the beta burst dynamics of cued voluntary movements in Tourette Syndrome"

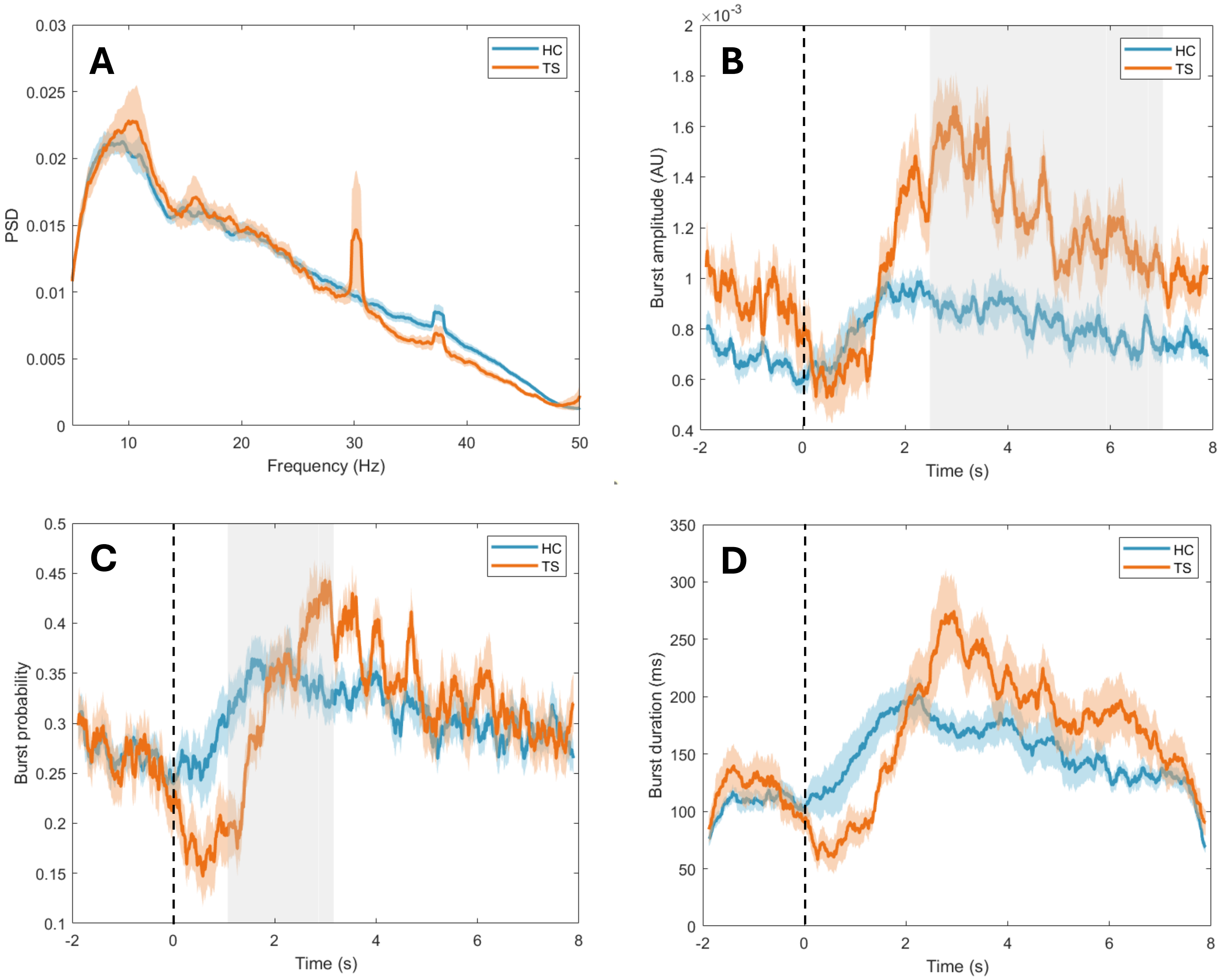


*Supplementary Figure 1. Trial averaged timecourses for characteristics of beta bursts within the contralateral primary motor cortex in individuals with Tourette Syndrome (orange) and neurotypical age- and sex-matched controls (blue). (A) power spectral density, (B) burst amplitude, (C) burst probability and (D) burst duration. A single index finger abduction and adduction occurred at time 0s. Data filtered 5-48Hz. Areas shaded in grey indicate a significant difference between the groups (p ≤ 0.05, corrected).*
